# Cellular Uptake and Fate of Cationic Polymer-Coated Nanodiamonds Delivering siRNA: A Mechanistic Study

**DOI:** 10.1101/2023.11.02.564900

**Authors:** Jan Majer, Marek Kindermann, Dominik Pinkas, David Chvatil, Petr Cigler, Lenka Libusova

## Abstract

Gene silencing using small interfering RNAs (siRNAs) is a selective and promising approach for treatment of numerous diseases. However, broad applications of siRNAs are compromised by their low stability in a biological environment and limited ability to penetrate cells. Nanodiamonds (NDs) coated with cationic polymers can enable cellular delivery of siRNAs. Recently, we developed a new type of ND coating based on a random copolymer consisting of (2-dimethylaminoethyl) methacrylate (DMAEMA) and N-(2-hydroxypropyl) methacrylamide (HPMA) monomers. These hybrid ND-polymer particles (Cop^+^-FND) provide near-infrared fluorescence, form stable complexes with siRNA in serum, show low toxicity, and effectively deliver siRNA into cells *in vitro* and *in vivo*. Here, we present data on the mechanism of cellular uptake and cell trafficking of Cop^+^-FND:siRNA complexes and their ability to selectively suppress mRNA levels, as well as their cytotoxicity, viability and colloidal stability. We identified clathrin-mediated endocytosis as the predominant entry mechanism for Cop^+^-FND:siRNA into U-2 OS human bone osteosarcoma cells, with a substantial fraction of Cop^+^-FND:siRNA following the lysosome pathway. Cop^+^-FND:siRNA potently inhibited the target GAPDH gene with negligible toxicity and sufficient colloidal stability. Based on our results, we suggest that Cop^+^-FND:siRNA can serve as a suitable *in vivo* delivery system for siRNA.

## INTRODUCTION

Over the past two decades, the enormous therapeutic potential of RNA interference has been recognized, resulting in the first regulatory approval of a small interfering RNA (siRNA)-based drug in 2018.^1^ The success of this therapeutic was based, among other components, on a lipid nanoparticle system designed to deliver the siRNA. Nevertheless, the inefficient penetration of siRNAs through cell membranes and their instability in a biological environment remain major obstacles to the broad therapeutic application of these molecules. Thus, alternative delivery/transfection systems with low toxicity and high protective capacity remain of utmost interest. Systems based on new types of lipid nanoparticles,^2,3^ polymers and polymer hybrids,^4^ dendrimers,^5,6^ inorganic nanoparticles and other materials have been investigated extensively.^2,7^

Nanodiamonds (NDs) are a promising platform for siRNA delivery because of their very low toxicity, material scalability and advanced surface modification possibilities. Efficient intracellular delivery of siRNA requires cationic NDs, which can form electrostatic complexes with the negatively charged siRNA. The siRNA molecules released from the ND surface then interfere with the expression of the target gene.^8^ The dissociation kinetics depend on the ND coating properties^8^ and can be tailored to meet specific demands for individual applications. A positively charged surface can be achieved either by hydrogenation^9–11^or surface modification of the ND with cationic moieties.^12^ Two types of NDs with different origins have been evaluated as siRNA transfection systems: detonation NDs (DNDs) and high pressure high temperature (HPHT) NDs. The characteristic advantage of DNDs is the small size of their primary crystals (a few nanometers), because the material can be excreted from a mammalian body through the renal pathway.^13^ DNDs modified with cationic polymers such as poly(ethyleneimine)^14,15^ and poly(amidoamine)^16^ and small cationic molecules such as ethylenediamine^17^ and lysine^18,19^ have been developed as siRNA vectors. In comparison with DNDs, HPHT NDs offer a unique advantage: they accommodate crystal lattice defects called nitrogen-vacancy centers that fluoresce in the near-infrared region.^20^ This fluorescence is extremely photostable and enables long-term tracking of cells,^21–23^ even in a single-particle model.^24^ Moreover, nitrogen-vacancy centers enable various optical quantum sensing modalities.^25–27^ Importantly, HPHT NDs modified with cationic polymers such as poly(ethyleneimine)^15,28–31^ and poly(allylamine)^15^ can deliver siRNA or microRNA to cells comparably to DNDs.

Nevertheless, the polymers used for coating of NDs are often relatively toxic and, after complexation with siRNA, do not stabilize the particles sufficiently in biological liquids. Recently, we focused on overcoming these limitations by designing a new type of ND copolymer coating based on (2-dimethylaminoethyl) methacrylate (DMAEMA). DMAEMA homopolymers and various DMAEMA-containing copolymers may serve as effective vectors for siRNA delivery.^32^ DNDs coated with DMAEMA grown from the surface using ATRP polymerization have been employed to deliver a DNA plasmid.^33^ To “dilute” the positive charge of the copolymer,^34^ we included N-(2-hydroxypropyl) methacrylamide (HPMA) monomers in the copolymer structure.^35,36^ NDs coated with this copolymer (Cop^+^-FND) were effective for siRNA transfection *in vitro* and *in vivo*. They also exhibited sufficient colloidal stability in serum,^35^ which makes them a promising candidate for further development of transfection agents.

Here, we study the mechanism of cellular uptake and cell trafficking of Cop^+^-FND complexes with siRNA (Figure 1A). We investigate the route of entry of these Cop^+^-FND:siRNA complexes using the specific inhibitors Dynasore and filipin. To assess and quantify changes in Cop^+^-FND:siRNA uptake in treated cells and controls, we exploit the bright and stable fluorescence of the FNDs. In addition to these mechanistic studies, we analyze the colloidal properties of the particles using dynamic light scattering (DLS) and electrophoretic light scattering (ELS) and investigate their cytotoxicity and viability. Finally, we quantify their ability to selectively suppress mRNA levels of glyceraldehyde 3-phosphate dehydrogenase (GAPDH) in cells.

**Figure 1.**
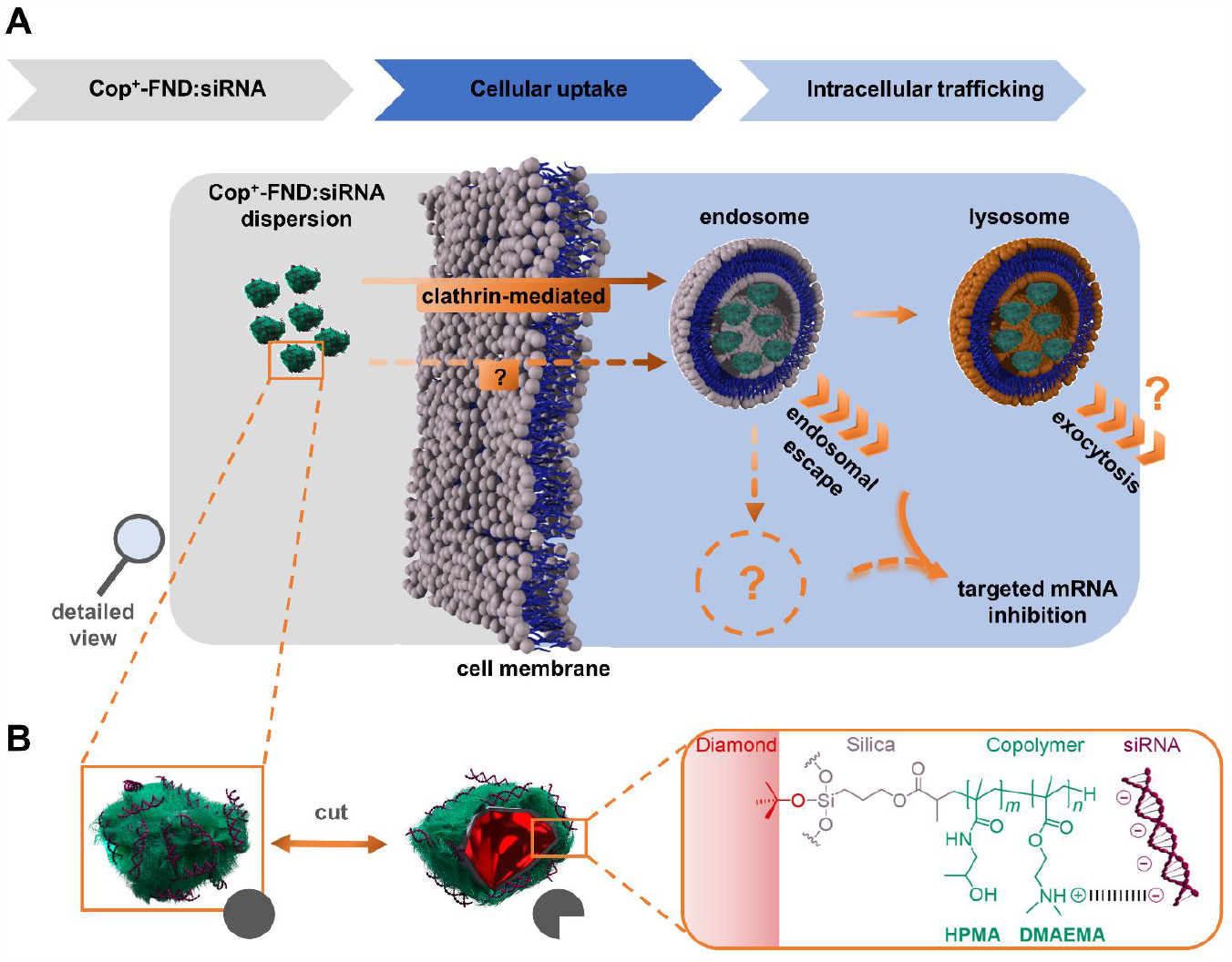
(A) Uptake routes and possible intracellular fates of Cop^+^-FND:siRNA complexes and (B) their schematic structure.

## MATERIALS AND METHODS

### Materials

Dimethylsulfoxide (DMSO), phosphate buffered saline (PBS) for cell culture, filipin complex from *Streptomyces filipinensis*, Dynasore, Triton X-100, Mowiol 4.88, GAPDH and β-actin primers, and DAPI were purchased from Merck (USA). Trypsin was purchased from Cytiva (USA). Dulbecco’s Modified Eagle Medium (DMEM) for cell culture with D/glucose, DMEM for cell culture with GlutaMAX, pyruvate, phenol red, D-glucose, fetal bovine serum (FBS) and Alexa 594 conjugated transferrin were obtained from Life Technologies™ (USA). RNeasy Mini Kit was obtained from Qiagen (Netherlands). Penicillin/streptomycin was purchased from Biowest (France). Glutamine, Lipofectamine^®^ RNAiMAX reagent, SuperScript III™ Reverse Transcriptase and paclitaxel were purchased from Thermo Fisher Scientific (USA). X-treme Gene HP transfection reagent, Cell Proliferation Reagent WST-1 and Cytotoxicity Detection Kit^PLUS^ (LDH) were obtained from Roche (Germany). 5× HOT FIREPol^®^ EvaGreen^®^ qPCR Supermix was acquired from Solis BioDyne (Estonia). Anti-GAPDH siRNA (siGAPDH, MW = 13,285) (sense strand: 5’-GGU CAU CCA UGA CAA CUU U(dT)(dT)-3’; antisense strand: 5’-AAA GUU GUC AUG GAU GAC C(dT)(dT)-3’), Atto425-labelled anti-GAPDH siRNA (siGAPDH-A425) and non-targeting siRNA (siCtrl, MW = 13,254) (sense strand: 5’-UAA GGC UAU GAA GAG AUA C(dT)(dT)-3’; antisense strand: 5’-GUA UCU CUU CAU AGC CUU A(dT)(dT)-3’) were synthesized by Sigma Aldrich. Early Endosome Antigen 1 (EEA1) specific rabbit monoclonal antibody (cat. no. mAb #3288) and Lysosome Associated Membrane Protein 1 (LAMP1) specific rabbit monoclonal antibody (cat. no. mAb #9091) were purchased from Cell Signaling Technology (USA). Anti-caveolin-1 rabbit polyclonal antibody was acquired from BD Biosciences (USA, cat. no. 610059). Secondary antibody Alexa Fluor^®^ 488 conjugated anti-rabbit donkey IgG was purchased from ThermoFisher Scientific (USA, cat. no. 711-545-152). All chemicals used for FND and Cop^+^-FND preparation have been previously described^36^.

### Cop^+^-FND preparation

High-pressure high-temperature diamond nanocrystals (Microdiamond, Switzerland, MSY 0–0.05 μm, type Ib with nitrogen impurity concentration ∼200 ppm) were oxidized by air in a furnace at 510 °C for 3 h followed by treatment in a mixture of H_2_SO_4_ and HNO_3_ (1:3) at 160 °C for 3 days. They were then washed with water, 1 M NaOH, 1 M HCl and water (centrifugation parameters have been described previously).^37^ To create NV centers in the diamond lattice, we modified the published protocol.^38^ Briefly, ND powder was irradiated with a 16.6 MeV electron beam followed by annealing (900 °C, 1 h) and air oxidation (510 °C, 3 h). The resulting powder (FNDs) was again treated with a mixture of H_2_SO_4_ and HNO_3_, washed with NaOH, HCl and water and then freeze-dried. This procedure was repeated once (sample irradiation and acid treatment). FND powder was redispersed in Milli-Q water (30 mL, 2 mg mL^−1^) by probe sonication as previously described.^36^ The resulting transparent colloid (ND) was incubated 30 min at room temperature and filtered using a 0.2 μm PVDF filter. FND dispersion was used for coating with a methacrylate-terminated thin silica layer using a modified Störber procedure;^39^ the amount of all components needed for silication procedure was linearly recalculated to the amount of FND powder (60 mg).^36^ The terminal methacrylate groups underwent radical polymerization, resulting in a dense layer of random copolymer poly{(2-dimethyl-aminoethyl methacrylate)-*co*-[N-(2-hydroxypropyl)methacrylamide]} (poly(DMAEMA-co-HPMA)). Detailed preparation (including purification) and characterization of FND@silica@poly[DMAEMA^+^-*co*-HPMA^0^] complexes was described previously^36^ (specifically, the sample denoted as “80^+^/20^0^” was used in this study). Briefly, HPMA and 2,2′-azobis(2-methylpropionitrile) (AIBN) were freshly recrystallized prior to use. Both DMAEMA (984 mg) and HPMA (328 mg) monomers were dissolved in DMSO (3.75 mL). AIBN (376 mg) was added, and the mixture was filtered using a 0.2 μm PTFE filter. Methacrylate-terminated FNDs in DMSO (376 μL) were added and the stirred mixture was secured in argon. The polymerization proceeded for 3 days under an argon atmosphere at 55 °C. The reaction was terminated by addition of MeOH. The resulting Cop^+^-FND samples were purified by centrifugation in nuclease free water to a final concentration of 4.1 mg mL^−1^ and stored at 4 °C.

*Bright-Field Transmission Electron Microscopy (TEM)* was performed with a JEOL JEM-1011 electron microscope operated at 80 kV equipped with a Tengra bottom-mounted camera as described previously.^37^ The TEM micrograph of the Cop^+^-FND is presented in Figure S8 in ESI showing a characteristic shape and size distribution of the FND particles (the polymer layer is not visible). Note the presence of individual separated particles which show no aggregation.

^*1*^*H NMR spectrum* of the Cop^+^-FND was recorded on a Bruker Avance III 500 spectrometer (499.88 MHz) equipped with a 5 mm PFG cryoprobe; 5 mg of ND sample was centrifugated 3x and transferred in D_2_O resulting in final volume approximately 50 μL. The signals were assigned by using a combination of 1D and 2D (H,H-COSY and H,C-HSQC) techniques. ^1^H NMR (400 MHz, D_2_O, δ) 4.02 (-OC**H**_**2**_-), 3.73 (-C**H**(OH)(CH_3_)), 2.90 (-NHC**H**_**2**_-), 2.62 (-OCH_2_C**H**_**2**_-), 2.20 (-N(C**H**_**3**_)_2_), 1.81 (C-C**H**_**2**_-, backbone), 1.02 (-CH(OH)C**H**_**3**_), 0.78 (C-C**H**_**3**_, backbone). The copolymer composition was estimated from ratios of the integral intensities of signals of DMAEMA^+^ (-OC**H**_**2**_-group) and HPMA^0^ (-C**H**(OH)(CH_3_) group).^36^

#### X-ray photoelectron spectroscopy (XPS)

XPS spectra were recorded using an Omicron Nanotechnology instrument equipped with a monochromatized AlKα source (1486.7 eV) and a hemispherical analyzer operating in a constant analyzer energy mode with a multichannel detector. A CasaXPS program was used for spectral analysis. The binding energy of C 1s (284.8 eV) was used for calibration of the binding energy axis.

### High-angle annular dark-field scanning transmission electron microscopy (HAADF STEM) with energy dispersive X-ray spectroscopy (EDS)

Samples (0.01 mg mL^−1^, 15 μL) were deposited on Quantifoil 2-1 holey carbon film coated copper TEM grids for 10 min. The remaining liquid was wiped up with a dust-free wipe (Kimwipe) and the grid was air-dried. STEM and EDS data acquisition was done on Jeol JEM-F200 (S)TEM operated at 200 kV. Images were acquired using HAADF detector with detecting angle 45-165 mrad at nominal camera length set to 150 mm. EDS data were acquired using JED 2300 X-ray spectrometer with a single 100 mm^2^ (0.98 sr) windowless SDD detector. Probe current was set to 0.85 pA and acquisition time for each area was 900 s. Presented elemental maps were calculated from the raw spectra using standardless Zeta factor method embedded in Jeol AnalysisStation software.

### Complexation of Cop^+^-FND:siRNA and Lipofect:siRNA

All components were redispersed in nuclease free water prior to use. By changing the amount of Cop^+^-FND at a fixed concentration of siGAPDH, we optimized the binding mass ratio for Cop^+^-FND:siGAPDH as previously described.^35,36^ We evaluated the siGAPDH binding efficacy and the apparent ζ-potential while keeping the Cop^+^-FND:siGAPDH mass ratio as low as possible (data not shown). We determined an optimal binding mass ratio of 26:1, which was used in all subsequent experiments. Cop^+^-FND:siGAPDH in a 26:1 mass ratio was prepared from siRNA (0.506 μL, 100 μM, 1.33 mg mL^−1^) diluted in Milli-Q water (4.28 μL). The diluted siRNA mixture was placed in a sonication bath (Bandelin Sonorex™ RK 31, 30/240 W, 35 kHz), and Cop^+^-FND (4.28 μL, 4.1 mg mL^−1^) was added in one portion. The resulting dispersion was sonicated in a sonication hotspot for and additional 30 s after mixing. The resulting transparent colloidal solution of Cop^+^-FND:siRNA complex was used in further experiments. Lipofect:siRNA lipoplexes were prepared by 5 min incubation of a mixture of Lipofectamine^®^ RNAiMAX reagent (3 μL) and siRNA (0.5 μL, 100 μM), both dissolved in serum-free DMEM medium.

### Assessment of colloidal stability by dynamic light scattering (DLS) and electrophoretic light scattering (ELS)

DLS intensity weighted diameter (Z-average) and average values of the apparent ζ-potential were obtained on a Zetasizer Nano ZSP (Malvern Instruments) equipped with a He–Ne laser (633 nm, 10 mW). Z-average diameters and polydispersity indexes (PDI) were inferred from the second-order time intensity autocorrelation function *g*^(2)^(*τ*)*-*1 considering 3^rd^ order cumulant analysis in Zetasizer Software 7.13. Unless stated otherwise, each sample was measured three times (unless stated otherwise) with an automatic duration at a backscatter angle of 173°; reported sizes represent a mean value of these measurements. The viscosities of serum-free DMEM and cell culture media (DMEM/10 % FBS/antibiotics) at 37 °C were set at 0.726^40^ and 0.738 cP,^40^ respectively. The dispersant refractive index was set at 1.33 for all solvents tested. ELS with a phase analysis of the scattered light at a 13° angle was used to determine the apparent ζ-potential. The samples were measured three times in Milli-Q water using a disposable cuvette (final volume 0.6 mL) with a dip cell and considering General purpose analysis (20-100 subruns). To calculate the apparent ζ-potential value from Henry’s equation, we used the Smoluchowski approximation for spherical noncoated particles without inspecting the value of the ratio of particle size to Debye length. For both types of measurements, the particle concentration was below 0.4 mg mL^−1^; dilution of the solvent by the testing sample was below 10%.

Supernatant collected from HeLa cells after incubation with Cop^+^-FND:siGAPDH complexes at different time points was measured without further dilution.

### Estimation of average number of siRNA duplexes per Cop^+^-FND particle

The count-based (molar) concentration of the Cop^+^-FND sample was estimated using nanoparticle tracking analysis (NTA). Nanosight NS300 equipped with a low-volume cell connected to a linear pump and 405 nm laser (<70 mW) was used. Stock solution of Cop^+^-FND sample (0.1 mg mL^−1^) was sonicated in the cup horn and diluted in water with dilution factor 2.5 × 10^2^ resulting in measurements with ∼45 particles per frame and number of valid tracks >7000. Camera level was set to 12 (shutter/gain: 1200/146), detected threshold to level 3 and syringe pump flow rate to level 120; blur, maximum jump distance, and minimum expected particle size were set to “auto”. The sample was measured 8 × 180 s at ∼25 °C. The concentration of siRNA was measured using Qubit fluorimeter with Qubit™ miRNA assay kit. The average number of siRNA duplexes per Cop^+^-FND particle was estimated from the molar concentration of the Cop^+^-FND (measured by NTA) and stock concentration of siRNA (measured by Qubit assay). The presence of free siRNA, i.e. <0.02 % of the total siRNA amount, was neglected (see the paragraph Release of siRNA from Cop^+^-FND:siRNA complex at different pH conditions in ESI).

### Cell culture

U-2 OS human bone osteosarcoma cells and HeLa human cervical adenocarcinoma cells were obtained from American Type Culture Collection and were cultured at 37 °C in a 5% CO_2_ atmosphere. Cells for qPCR experiments were cultivated in DMEM medium supplemented with 10% FBS in 1% penicillin/streptomycin and passaged every 3 days. Cells for imaging experiments were cultured in a phenol red-free medium.

### RNA interference evaluation

U-2 OS wildtype cells were seeded in 12-well plates (0.5 mL cell culture media). Confluent cells (70%) were transfected with Cop^+^-FND:siGAPDH and Cop^+^-FND:siCtrl (diamond@copolymer:siGAPDH mass ratio of 26:1) to final concentrations of 35 μg mL^−1^ Cop^+^-FND and 100 nM siRNA. Furthermore, cells were transfected with Lipofect:siRNA (siGAPDH, siCtrl) lipoplexes to final concentration of 100 nM siRNA. After 36 h of incubation, cells were washed with PBS and harvested. Total RNA was extracted from cells using the RNeasy Mini Kit. A 500 ng aliquot of total RNA was transcribed to cDNA with SuperScriptIII^®^ reverse transcriptase, according to the manufacturer’s protocol. Reverse-transcribed cDNA was diluted 20×. HOT FIREPol^®^ EvaGreen^®^ Supermix was used for quantitative real-time polymerase chain reaction (qPCR) analysis. Each polymerase reaction was performed in 15 μL of a reaction mixture containing 3 μL HOT FIREPol^®^ EvaGreen^®^ Supermix (5×), 0.6 μL GAPDH and β-actin primers (Sigma Aldrich), 6.4 μL H_2_O and 5 μL cDNA. An IQ5 Multicolor RT PCR Detection System (BioRad, USA) was used for gene expression analysis. The comparative threshold (CT) was analyzed. The quantitative qPCR data were determined relative to untreated cell samples. Using the ΔΔCT method,^41^ GAPDH gene expression levels were normalized to expression levels of β-actin and compared to treatment with non-targeting siCtrl.

### Cell viability and cytotoxicity assay

*In vitro* cytotoxicity of the complexes was evaluated using U-2 OS cells and an LDH assay kit.^42^ Cells were counted with a Countess II Automated Cell Counter (ThermoFisher Scientific, USA) and seeded in 96-well plates at a density of 10,000 cells per well in 100 μL of full DMEM. Cells were treated either with Cop^+^-FND:siRNA (50.6 μg mL^−1^: 1.92 μg mL^−1^/145 nM and 202 μg mL^−1^: 7.68 μg mL^−1^/578 nM) or Lipofect:siRNA (0.84 % v/v: 140 nM). After 36 h, negative control cell samples were treated with Triton X-100 (4%, v/v). After 10 min, 100 μL of supernatant from samples was transferred to new wells in a 96-well plate. LDH catalyst (2.17 μL) and LDH dye (97.8 μL) were added to the supernatant samples and incubated (15 min, 37 °C, 5% CO_2_). Finally, a stop buffer (50 μL) was added. The WST-1 assay was used to examine cell proliferation.^42^ Samples were seeded and transfected as in the LDH assay. Moreover, negative controls were incubated with paclitaxel (40 nM). After 36 h, WST-1 reagent (10 μL) was added and incubated for 45 min. Finally, cells were examined colorimetrically with an ELx800 plate reader (Dynex, Czech Republic) (emission 450 nm/background 630 nm). Measurements from blank wells were used as a baseline for sample well measurements. All experiments were performed in triplicate.

### Cellular uptake rate

U-2 OS wildtype cells were plated on glass coverslips in 24-well plates (60% confluence, in 350 μL of cell culture media). The next day, fresh aliquots of Dynasore and filipin in DMSO were prepared. Cells were pre-treated with Dynasore or filipin at final concentrations of 400 μM or 5 μg mL^−1^,^43^ respectively. The final concentration of DMSO in all samples was 2% v/v. After 20 min, cells were chilled (3 min, 7 °C). Cop^+^-FND:siRNA complexes were pipetted into sample wells (5 min, 7 °C, 10 % FBS) with a resulting concentration of 50.6 μg mL^−1^ Cop^+^-FND and 1.92 μg mL^−1^/144.6 nM siRNA. To prevent potential aggregation of the particles, the cell culture was cooled down prior to transfection. Afterward, the cells were co-incubated with the complexes for 55 min at 37 °C. Finally, cells were washed with PBS and fixed with paraformaldehyde (3%) in PBS (15 min). Cells were permeabilized with Triton X-100 (1%, v/v) and blocked with 2% bovine serum albumin (BSA) for 5 min. All antibodies were diluted in 2% BSA dissolved in PBS. Cells were stained with primary antibody against Caveolin-1 (Cav-1), EEA1 or LAMP1 for 90 min followed by incubation with the secondary antibody conjugated with Alexa Fluor 488 for 60 min. Images were acquired using a Zeiss LSM 880 confocal microscope (Carl Zeiss AG, Germany). The intracellular area of Cop^+^-FND:siRNA complexes was quantified by the “Analyze particle…” tool in Fiji ImageJ^44^ with the following parameters: Size (micron^2): 4–inf, Circularity: 0.3–1.0, Pixel units: Yes.

### Intracellular Cop^+^-FND:siRNA trafficking

U-2 OS wild-type and polyclonal U-2 OS expressing plasmids pcDNA_PGK_Hygro-GFP-Rab5A (U-2 OS Rab5) and pcDNA_PGK_Hygro-GFP-Rab7A were seeded on coverslips in 24-well plates (60% confluence). Both GFP-Rab5A and GFP-Rab7 were subcloned from the original plasmids (Addgene #49888 and GFP-Rab7^45^) into pcDNA3.1 Hygro vector backbone (cat. no. V87020, ThermoFisher Scientific, USA). To reach nearly physiological levels of expression, the expression was driven from a PGK promoter. The next day, cells were transfected with Cop^+^-FND:siRNA complexes (50.6 μg mL^−1^: 1.92 μg mL^−1^/145 nM) in full DMEM without phenol red. After 6 h, cells were washed with PBS, fixed and mounted on glass slides in Mowiol at multiple time points (0–360 min). Moreover, wild-type cells were permeabilized by Triton X-100 (0.1%) and stained with anti-EEA1 or anti-LAMP1 antibody followed by secondary antibody staining and sample mounting in Mowiol (10% w/v). Samples were captured with a Zeiss LSM 880 confocal microscope. Experiments were conducted in duplicates and triplicates. Colocalization was expressed by Pearson’s correlation coefficient (*PCC*) calculated in Fiji ImageJ (Coloc2 plugin) performing one-to-one pixel matching analysis to evaluate spatial correlation. Regions of interest (ROIs) – cell boundaries – were drawn manually using Fiji ImageJ based on green fluorescent protein (GFP), A488 and autofluorescence signal.

### Confocal microscopy setup

Image data were acquired with a Zeiss LSM 880 Axioobserver laser scanning confocal microscope equipped with lasers emitting power of 30 (attenuator set to 1% of maximum intensity), 25 (5%) and 20 (7–10%) mW at wavelengths of 405, 488 and 561 nm, respectively. The microscope was controlled by ZEN Black software. An oil immersion objective (63×, 1.4 NA) was used. Images of 976 × 976 pixels were acquired at 70 nm pixel size and 8.83 μm pixel dwell time. Acquisition parameters were identical for each fluorophore. Lambda scans – series of individual images at set wavelength range of detection – were obtained during imaging to separate spectral features of spectrally overlapping components through linear unmixing feature (ZEN Black, Zeiss).

## RESULTS AND DISCUSSION

### Cop^+^-FND forms colloidally stable complexes with siRNA

The architecture of our vectors is based on a random cationic copolymer interface covalently grafted from a silica-coated FND core (Cop^+^-FND).^35,36^ The charge-diluted copolymer layer contains the cationic comonomer DMAEMA (80% w/w, final percent mass in copolymer layer),^36^ allowing electrostatic interactions with negatively charged siRNA. The electroneutral comonomer HPMA (20% w/w) “dilutes” the positive charge. To graft the polymer onto the particles, we used the so-called “grafting through” approach.^46,47^. In this approach, the polymerizable methacrylate groups are first anchored onto the nanoparticle surface, and radical polymerization is initiated in a solution containing both free monomers and methacryloylated nanoparticles. The pre-attached polymerizable moieties are gradually integrated into the growing polymer chains. “Grafting through” thus differs from the “grafting from” method, in which functional initiators are chemically linked onto the nanoparticle surface to directly initiate polymerization. Although the “grafting through” approach typically provides lower grafting densities and layer thicknesses than the “grafting from” approach, the amount of polymer attached is largely independent of the polymerization conditions. This polymerization approach was thus classified as robust, error-tolerant and reproducible.^48^

We used both monomers (DMAEMA, HPMA) in a mixture that provided Cop^+^-FND particles with a random composition of the copolymer (see Figure 1B). The expected cationic character of the copolymer required for interaction with the negatively charged siRNA was confirmed by ELS showing an apparent ζ-potential of 35.8 ± 3.9 mV in water (Table S1 in ESI). The particles were colloidally stable with corresponding Z-average diameter of 126 ± 1 nm (calculated from 2 DLS measurements). The estimated average number of siRNA duplexes per nanoparticle was ∼1360; this estimate is based on the NTA of the Cop^+^-FND sample that provides a count-based concentration.

The Cop^+^-FND:siGAPDH complex with an optimal mass ratio of 26:1 (see Experimental) was prepared by slowly adding Cop^+^-FND solution to siRNA solution under constant sonication in a bath. To confirm the binding of the siRNA to the copolymer layer, we analyzed elemental composition of the layer using HAADF STEM with energy dispersive X-ray spectroscopy (EDS). We focused on analysis of phosphorus content, because this element is solely present in siRNA, but not in the original polymer layer and X-ray signal of P was successfully used to detect even a single layer of nucleic acid on carbon based support.^49^

STEM was able to clearly visualize individual nanodiamond particles. In combination with an X-ray detector, we collected X-ray spectra from each pixel of a STEM image and calculated elemental maps based on the collected data. Figure 2 presents such maps showing single Cop^+^-FND and Cop^+^-FND:siGAPDH particles. The core diamond particles are carbon based and both of them are surrounded by a layer of polymer containing a significant percentage of nitrogen. While nitrogen is present in both polymer and siRNA, phosphorus is present only in the siRNA. Correspondingly, we observed no P signal in Cop^+^-FND (Fig. 2B and C) and a clear P signal colocalizing with the N signal in Cop^+^-FND:siGAPDH containing siRNA (Fig. 2A and C). We can thus confirm that siRNA binds to the polymer layer and remains bound in Cop^+^-FND:siGAPDH under the experimental conditions used.

**Figure 2.**
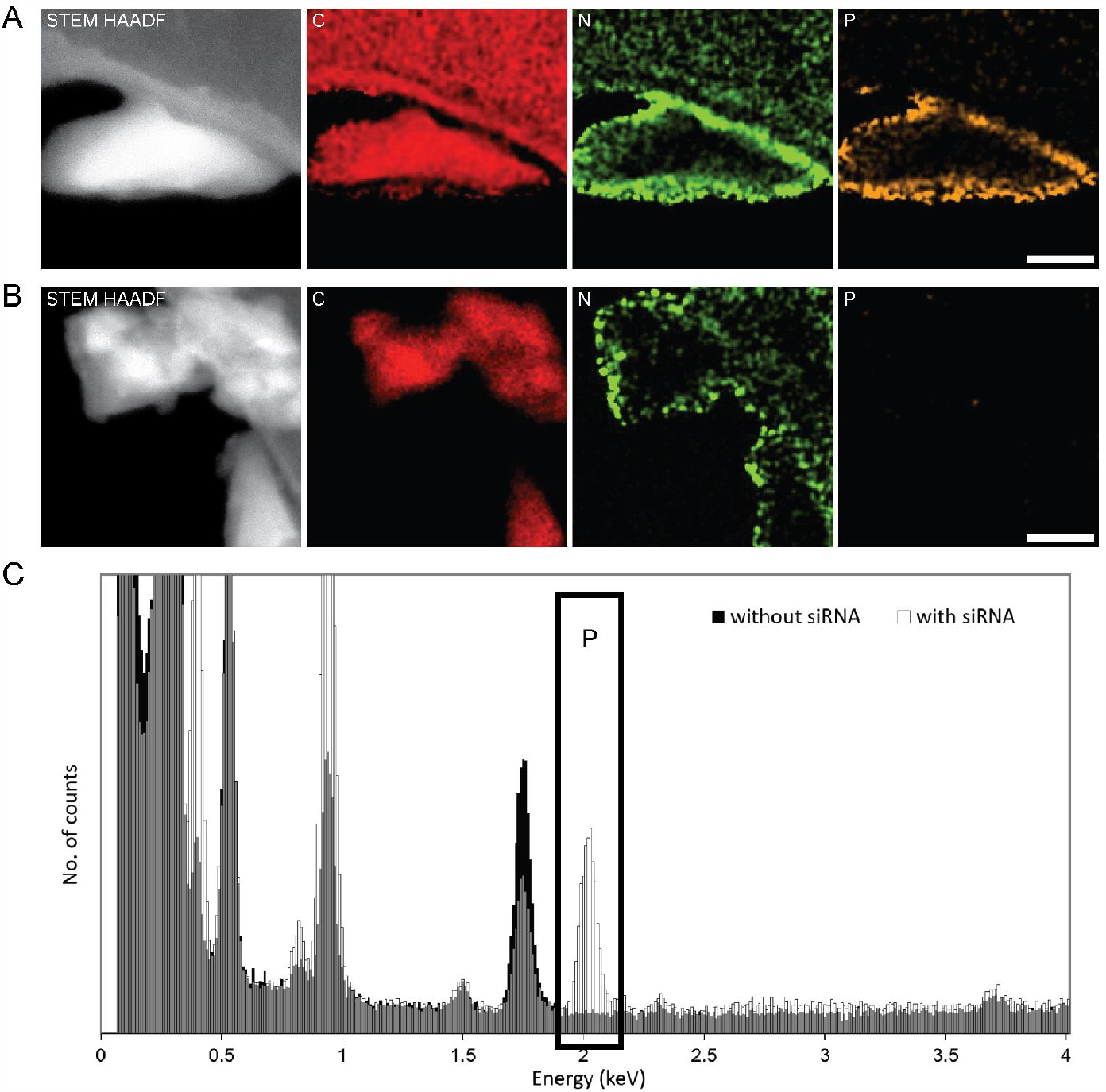
(S)TEM EDS: (A) and (B) show from left to right STEM-HAADF image intensity map C EDS signal (based on Kα peak at 0.277 keV) and quantitative maps of N and P for (A) Cop^+^-FND:siGAPDH and (B) Cop^+^-FND particles. Scalebar 25 nm. Graph in (C) shows comparison of overlaid X-ray spectra collected from both displayed particles. The Kα peak of P at 2.013 keV highlighted by a rectangle indicates a significant amount of P from siRNA present in Cop^+^-FND:siGAPDH in contrast with a flat background of Cop^+^-FND.

We also analyzed the elemental composition of the FND, Cop^+^-FND and Cop^+^-FND:siGAPDH using XPS. The obtained data (Figure S10 and Table S4 in ESI) correspond well with the EDS results. Most importantly, the phosphorus (0.21 at. %) was present only in the Cop^+^-FND:siGAPDH, documenting the successful formation of the complex with siRNA. Nitrogen was found in both Cop^+^-FND and Cop^+^-FND:siGAPDH, while the starting FND did not contain any measurable content of N. Interestingly, the binding of siRNA led to a measurable increase in nitrogen content from 3.5 at. % in Cop^+^-FND to 5.6 at. % in Cop^+^-FND:siGAPDH. Similarly, the oxygen concentration raised from 10 at. % in FND to 18 at. % in both Cop^+^-FND and Cop^+^-FND:siGAPDH. The Si content originating from the underlying silica layer was similar for both samples. By similar reasoning as above for EDS, we can conclude that the quantitative XPS results are consistent with the expected composition of both Cop^+^-FND and Cop^+^-FND:siGAPDH samples.

Next, we investigated the temporal stability of the optimized Cop^+^-FND:siGAPDH complex in full cell culture media at 37 °C. Monitoring of the complex stability over approximately 30 min revealed a slight broadening in size distribution and increase in mean size (for more detailed DLS data, see Figure S1 in ESI). To gain deeper insight into the colloidal behavior of Cop^+^-FND:siGAPDH *in vitro*, we assessed complex stability in a model experiment in full cell culture media at 37 °C after incubation with HeLa cells at different time points (0, 10, 20 and 40 min) (Figure 3A). We observed slow aggregation of the sample (typical DLS results are shown in Figure S1B). The cumulant-based polydispersity index and intensity-weighted mean size increased, while a decrease in the scattered intensity (after 40 min) was likely caused by a lower number of particles in the measured volume as a consequence of partial particle sedimentation.

**Figure 3.**
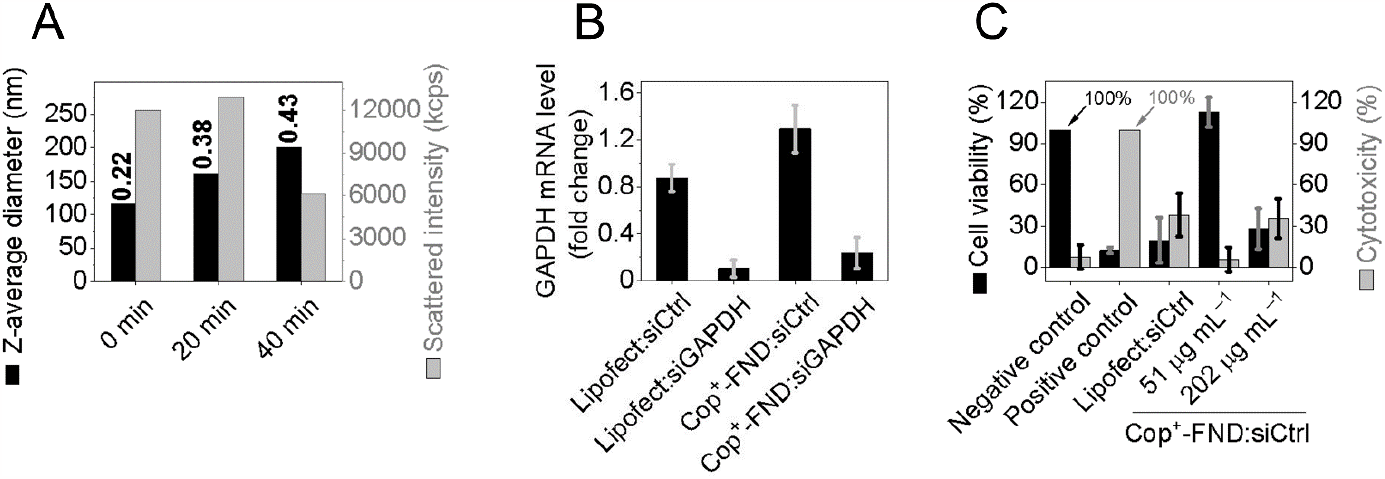
Stability, *in vitro* inhibition efficacy, viability and cytotoxicity of the Cop^+^-FND:siGAPDH complex (mass ratio 26:1). (A) Temporal dependence of the Z-average diameters (measured by DLS) and the corresponding scattered intensity signals measured after incubation with HeLa cells at 37 °C (supernatant was analyzed at 25 °C). The numbers above the Z-average bars indicate the cumulant-based polydispersity index (PDI). (B) Inhibition of GAPDH mRNA expression in U-2 OS cells measured by RT-qPCR, 36 h post-treatment with a fixed 100 nM concentration of siRNA (*n* = 3 experiments). A commercial transfection reagent Lipofectamine RNAiMAX was used as a negative or positive control (if complexed with siCtrl or siGAPDH, respectively). (C) Cytotoxicity in U-2 OS cells: LDH assay assessing the release of lactate dehydrogenase from damaged cells. Positive control represents cells treated with Triton X-100. Viability: WST-1 assay assessing the effect of Cop^+^-FND:siGAPDH on cellular metabolic activity. Positive control represents cells treated with paclitaxel. The experiments were performed at Cop^+^-FND concentrations of 51 and 202 μg mL^−1^ (*n* = 3 experiments). Negative control represents cells incubated in fresh culture medium (10% FBS). Data for panels B and C display mean ± sample standard deviation.

### Cop^+^-FND:siRNA inhibits GAPDH mRNA levels with low cytotoxicity

To evaluate the inhibition efficiency and toxicity of Cop^+^-FND:siRNA, U-2 OS human bone osteosarcoma cells were incubated with the complexes for 36 h (Figure 3B, C). We observed significant inhibition of GAPDH mRNA levels in the presence of 100 nM Cop^+^-FND:siGAPDH and a commercial transfection reagent RNAiMAX complexed with siGAPDH as positive control (Lipofect:siGAPDH; 100 nM siGAPDH) compared with their counterparts containing non-targeting siRNA (siCtrl). Cell viability and cytotoxicity remained unaffected even at a higher, 140 nM concentration of Cop^+^-FND:siCtrl (also see previous work).^35^ In contrast, addition of Lipofectamine RNAiMAX complexing 140 nM siCtrl (140 nM) greatly decreased the cell viability, down to ∼20%. To obtain a similarly low level of cell viability for Cop^+^-FND:siCtrl, a 4-fold higher concentration of the Cop^+^-FND was needed (560 nM siCtrl). Consistently, Cop^+^-FND concentrations of 560 nM led to increased cytotoxicity levels, similar to those measured in 140 nM Lipofect:siCtrl samples.

### Cop^+^-FND:siGAPDH enters cells primarily *via* clathrin-mediated endocytosis

NDs designated for active siRNA delivery need to adhere to the cell surface and undergo internalization, intracellular trafficking and endosomal release to the cytoplasm. There are four main mechanisms of cell entry for nanoparticle uptake: phagocytosis, macropinocytosis, clathrin-mediated endocytosis and caveolin-mediated endocytosis.^50^ Phagocytosis is most prominent in the cells of the immune system.^51^ However, its relevance for nanoparticle cell entry in non-professional phagocytes is unknown. The other three routes―all forms of endocytosis―are known to participate in FND internalization.^52–56^ Their uptake rates typically differ among cell types.^57–59^ Moreover, a protein corona, which is formed by adsorbed constituents of cell culture media onto the surface of FNDs, affects the nature of nanoparticle-cell interactions and the uptake mechanism.^60,61^ For cationic, poly(ethyleneimine)-coated FNDs bearing siRNA, macropinocytosis serves as the key internalization mechanism enabling the biological effect of the delivered siRNA.^28^

After internalization, NDs may be destined for degradation through lysosomal pathways. As the carbon lattice of the NDs is non-degradable by lysosomal hydrolytic machinery, FNDs may undergo exocytosis to the extracellular space.^62,63^ Alternatively, unmodified FNDs may escape into the cytoplasm by penetrating the endosomal membrane with their sharp edges.^52,64,65^ These sharp-shaped FNDs thus effectively avoid exocytosis by lysosomal fusion with the cell membrane. In contrast, rounded FNDs are hastily exocytosed from the cell, as they do not leave the endosome-lysosome-exosome route.^64^ This observation has important implications for drug/gene delivery because bound active compounds need to relocate to the cytoplasm to have a biological effect.^52^ Nevertheless, data on the intracellular fate of FNDs has only been documented for uncoated, anionic particles. The situation may be different for FNDs coated with cationic polymers.

To uncover the uptake mechanism of Cop^+^-FND:siGAPDH complexes, we used well-established specific inhibitors of two main endocytic pathways. Dynasore inhibits clathrin-mediated endocytosis by controlling the biological activity of dynamin GTPase, while filipin blocks caveolae-mediated endocytosis by interacting with membrane sterols and altering membrane permeability.

We pretreated cells with previously evaluated concentrations^43,66^ of these inhibitors for 30 min and then added 140 nM Cop^+^-FND:siGAPDH. After 60 min incubation, we fixed and analyzed the cells for the FND fluorescence. The fluorescence intensity coming from the internalized nanoparticles was similar in non-treated, mock-treated and filipin-treated cells (Figure 4; Figure S3 in ESI). This implies that the caveolae-mediated internalization pathway is not significantly involved in Cop^+^-FND:siGAPDH uptake.

**Figure 4.**
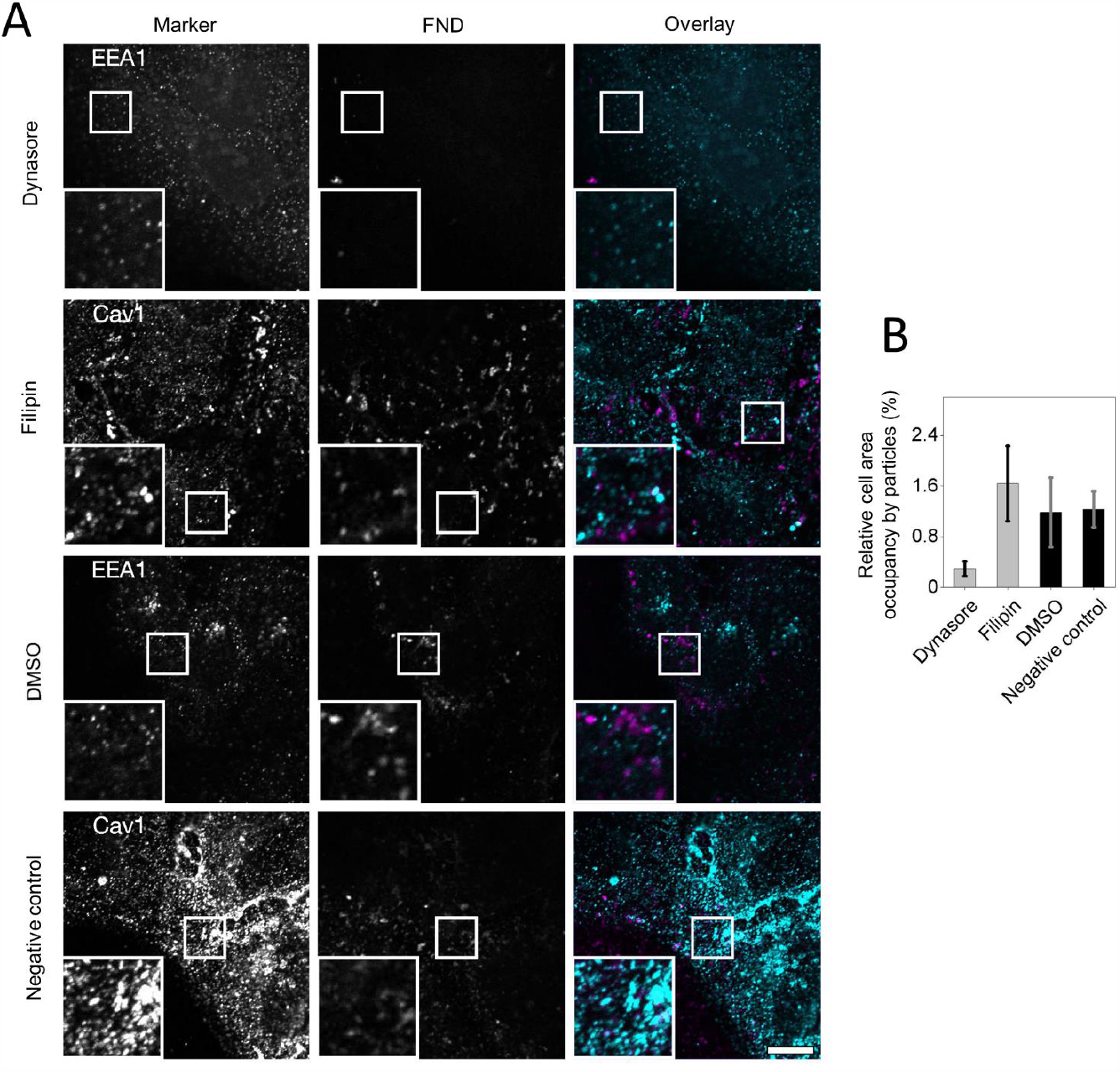
U-2 OS cells incubated in serum-free DMEM with filipin (5 mg mL^−1^) or Dynasore (400 μM) for 30 min and then with Cop^+^-FND:siGAPDH (140 nM, magenta) for an additional 60 min. Positive control cells were incubated with Cop^+^-FND:siGAPDH only. (A) Fixed cells treated with Dynasore and stained with anti-EEA1 antibody; fixed cells treated with filipin and stained with anti-Cav1 antibody. Both samples were then incubated with secondary anti-rabbit IgG-Alexa Fluor 488 (A488 Ab, cyan). The scale bar represents 20 μm. (B) Quantitative analysis of the inhibitory effects of Dynasore and filipin on Cop^+^-FND:siRNA uptake expressed as relative cell area occupancy by Cop^+^-FND:siGAPDH particles (*n* = 2 replicates).

On the other hand, the complex uptake was significantly decreased in the Dynasore-treated cells. First, using transferrin as a cargo control, we determined that Dynasore inhibits the clathrin-mediated endocytosis at 400 μM concentration (Figure S2; Figure S4 in ESI). Next, we tested the influence of the same Dynasore concentration on Cop^+^-FND:siGAPDH uptake. The overall fluorescence from FNDs decreased significantly compared to the signal from the positive control group (Figure 4; Figure S5 in ESI). This finding implies that in U-2 OS cells, the predominant entry mechanism for Cop^+^-FND:siGAPDH involves GTPase dynamin, which is essential for clathrin-mediated endocytosis. Indeed, clathrin-mediated endocytosis is considered the main internalization mechanism for NDs.^55,67^ However, Dynasore was not able to completely prevent the uptake of the complexes. Approximately one fourth of the fluorescence signal was still detectable in Dynasore-treated cells. We cannot exclude the possibility that the clathrin-mediated pathway was not fully inhibited, and the residual activity allowed delivery of a limited amount of Cop^+^-FND:siGAPDH. However, it is more likely that our cells used multiple uptake pathways, such as macropinocytosis or phagocytosis, as observed for other types of nanoparticles.^67–69^ Finally, Dynasore is known to modulate levels of cholesterol in the plasma membrane,^70^ which could also affect other uptake pathways.

### A substantial fraction of Cop^+^-FND:siGAPDH enters the lysosomal pathway

To understand the dynamics of Cop^+^-FND:siGAPDH inside cells, we analyzed their localization in a time series. First, we quantified particle uptake over 15-360 min using confocal microscopy (Figure 5A). Although endocytosis is a very rapid process in mammalian cells,^71^ we did not observe a significant signal from FNDs inside cells for the first 15 min of incubation. However, confocal microscopy proved that some particles were already present on the plasma membrane (data not shown). Subsequently, the intracellular signal from FNDs rose between 15 and 120 minutes and stayed fairly stable for the next few hours (Figure 5A).

**Figure 5.**
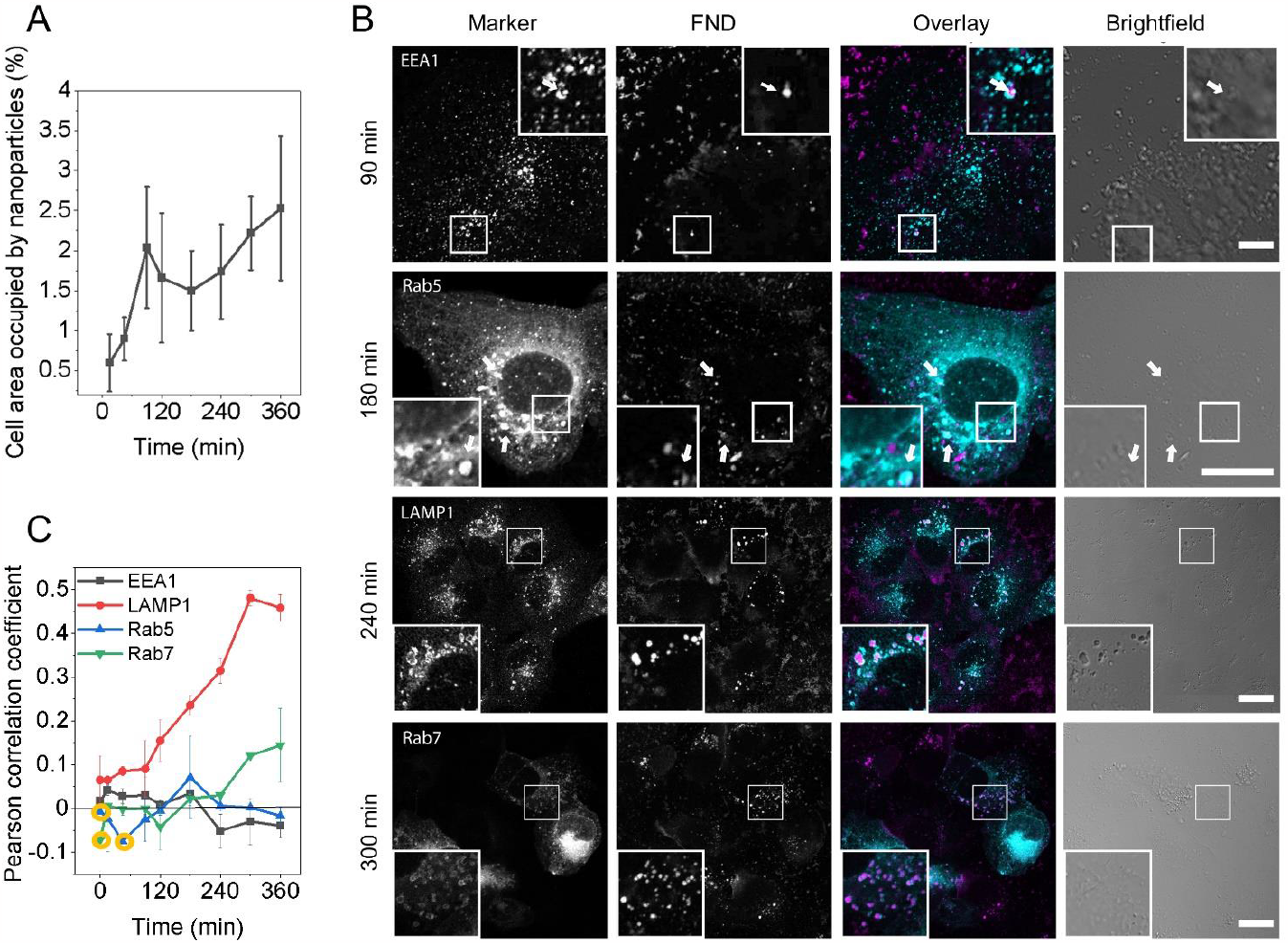
Temporal study analyzing colocalization of the FND signal with various endosomal structures and dynamics of complex uptake. U-2 OS cells were incubated with Cop^+^-FND:siGAPDH complexes for up to 6 h and then fixed in paraformaldehyde. (A) The gradual increase in the number of nanoparticles residing within the boundaries of U-2 OS cells in a single plane was observed by particle analysis (ImageJ). (B) Confocal images of i) wildtype cells (first and third row) immunostained with primary antibodies against EEA1 and LAMP1 followed by incubation with secondary antibody IgG-Alexa Fluor 488 (cyan) and ii) polyclonal cells (second and fourth row) expressing GFP-Rab5/Rab7 fusion proteins (cyan). Both cell cultures were incubated with Cop^+^-FND:siGAPDH complexes (140 nM) (magenta). Arrows exemplify overlaps between the vesicular fluorescent marker and the FND signal. The scale bars represent 20 μm. (C) Colocalization analysis expressed by Pearson correlation coefficient between FNDs and endosomal structures (*n* = 2 experiments). Points in orange circles represent time points at which only one sample was acquired.

Next, we studied the intracellular fate of Cop^+^-FND:siGAPDH using temporal colocalization studies of Cop^+^-FND:siGAPDH with several vesicle markers. To track early endosomes, we monitored both EEA1 and Rab5 proteins, which mark the same cell compartment and allowed us to corroborate the reproducibility of our microscopy observations (EEA1 protein is one of the key effectors of Rab5 GTPase protein).^72^ To assign late endosomes, we used Rab7 GTPase;^73^ for lysosomes, we assessed LAMP1 protein marker.^74^ Our U-2 OS cells expressed GFP fusions of Rab5 and Rab7, enabling their direct observation, while EEA1 and LAMP1 were immunostained with fluorescently labeled antibodies. We observed some complexes to localize in all compartments studied at some time point (Figure 5B). However, quantification of colocalization by the Pearson correlation coefficient did not reveal any significant accumulation in early endosomes labelled by EEA1 and Rab5 at any time point (Figure 5C). Rather, the particles accumulated over time in late endosomes and predominantly in growing lysosomes. This suggests that a substantial fraction of Cop^+^-FND:siGAPDH complexes was confined to early endosomes only for a short time and readily progressed toward lysosomes. The cell journey of Cop^+^-FND:siGAPDH (or Cop^+^-FND) then likely ended by exocytosis, which is a typical pathway for elimination of nanoparticles from cells.^75^ We observed ND fluorescent signals within the cells 15 min after introducing the nanoparticle complexes into the cell culture media, suggesting an early uptake startpoint. The uptake of particles increased steadily until 120 min. Thereafter, the intracellular area occupied by NDs changed only modestly until the 6 h timepoint (Figure 5A). The observed drop in ND area within a cell may be connected with the increased colocalization between NDs and lysosomes (Figure 5C, LAMP1), wherein the NDs would be tightly packed.

We note that internalized nanoparticles can also reach lysosomes via alternative routes, such as macropinocytosis or phagocytosis. While we were tracking the Cop^+^-FND:siGAPDH complexes associated with vesicles, there was also a second population of particles widely scattered throughout the cytoplasm that was not associated with any of our vesicular markers. The manner in which they entered the cell and their subsequent fate remain elusive. The observed strong silencing effect of siGAPDH (Figure 3B) was thus related either to the efficient escape of Cop^+^-FND:siGAPDH (or siGAPDH) from endosomes or from these unidentified vesicles.

### Studies on release of cargo siGAPDH from Cop^+^-FND

We hypothesized that successful delivery of fluorescently labeled siGAPDH to the cytosol followed by dissociation of siGAPDH would manifest in spatial separation of the FND and siGAPDH signals. We thus followed the spatiotemporal fate of Cop^+^-FND:siGAPDH in cells (Figure S6 in ESI). While we detected the FND signal soon after the start of incubation, bright regions corresponding to siGAPDH began to appear later. At the 90 min timepoint, we observed a small number of vesicles with an FND signal overlapping the siGAPDH signal. At later timepoints (240 and 300 min), the number of these vesicles increased, the diameter and the intensity of both FND and siGAPDH signals increased, and the vesicles were located in the presumed perinuclear region. Nanoparticles aggregated in large clusters inside cells were also observed in the brightfield, and the contrast increased with time. Importantly, the siGAPDH signal was absent in a substantial population of FND-containing vesicles, while some vesicles contained only a siGAPDH signal (see the arrows). This observation suggests siGAPDH releases from the nanoparticles; however, our imaging capabilities were not sufficient to document this process in detail and localize the release events.

The Pearson’s correlation coefficient of FND and siRNA showed an increasing trend of correlation between the two markers until a putative plateau was reached at the 5 h and 6 h time points (Figure S7 in ESI). We attribute the increasing trend to the increase of fluorescence signal intensity above the limit of detection, which occurred predominantly during the nanoparticle accumulation in lysosomes. Moreover, the Pearson’s correlation coefficient at time 0 min and undosed (control) cell samples showed high variability, which was reflected in the large standard deviation. The analysis error decreased with increasing marker fluorescence signal inside the cells. Further, we attempted to determine the released fraction of siRNA in pH range ∼4-7 using quantitative fluorescence-based assay (Tables S2 and S3 and the accompanying text in ESI). While we confirmed the above investigated formation of stable complexes of Cop^+^-FND with siRNA, we did not observe a measurable release of siRNA from Cop^+^-FND:siGAPDH at any pH. Nevertheless, due to the observed silencing activity of siGAPDH, which can only proceed in cytoplasm, at least some fraction of siGAPDH reached the cytoplasm in a molecular form. This conclusion is supported by the work of Xu *et al*., who reported that electrostatic binding of siGAPDH and cationic nanoparticles is most likely disrupted at neutral pH in combination with competitively binding cationic proteins, suggesting the likelihood of siGAPDH separating at either an early stage of endosome entrapment or within the cytoplasm.^11^ Therefore, it is likely that a fraction of siRNA that was below the limit of detection escaped by the endosomal pathway.

## CONCLUSIONS

Cop^+^-FND:siRNA complexes are recently developed transfection nanosystems that show promising stability and biological efficiency for *in vitro* and *in vivo* delivery of siRNA. Previous studies of Cop^+^-FND:siRNA complexes have focused mainly on their physicochemical properties^36^ and bioavailability/distribution after administration to the body.^35^ Herein, we investigated the uptake mechanism and intracellular fate of these complexes to provide more comprehensive knowledge about this delivery platform. To visualize the particles, we exploited the extremely photostable NV centers in NDs. We found that the predominant (>70%) entry mechanism for Cop^+^-FND:siRNA is clathrin-mediated endocytosis. Although a significant fraction of Cop^+^-FND:siRNA followed the lysosome pathway, which typically weakens the siRNA silencing effect, we observed effective transport of siRNA into U-2 OS human bone osteosarcoma cells, leading to a remarkable inhibition of targeted GAPDH mRNA. The efficacy of Cop^+^-FND:siRNA was fully comparable to the performance of commercial Lipofectamine RNAiMAX transfection agent. Based on our results, we suggest that U-2 OS cells likely combine multiple uptake routes, as shown for uptake of other types of nanoparticles into mammalian cells.^67–69^ In our microscopy study, we also observed a gradual dissociation of the fluorescently labeled siRNA from the Cop^+^-FND:siRNA complexes, which is crucial for the observed silencing effect. This effect was likely not caused just by pH, as we determined in a release experiment at various pHs. Finally, we note that we did not focus on Cop^+^-FND:siRNA exocytosis in this work, which would deserve a specialized separate study. Overall, we consider Cop^+^-FND:siRNA to be promising delivery system for siRNA that deserves further exploitation as an *in vivo* therapeutic nanosystem. Animal studies are currently underway.

## Author Contributions

Jan Majer: Formal Analysis, Investigation, Methodology, Validation, Visualization, Writing – original draft, Writing – review & editing. Marek Kindermann: Formal Analysis, Investigation, Methodology, Validation, Visualization, Writing – original draft, Writing – review & editing. David Chvatil: Investigation, Writing – original draft. Petr Cigler: Conceptualization, Formal Analysis, Funding acquisition, Methodology, Project administration, Supervision, Validation, Writing – original draft, Writing – review & editing. Lenka Libusova: Conceptualization, Formal Analysis, Funding acquisition, Methodology, Project administration, Supervision, Validation, Writing – original draft, Writing – review & editing.

## Acknowledgements

The authors are grateful to Vaclav Bocan (Faculty of Science, Charles University) for help with confocal microscopy and to Jakub Copak (Institute of Organic Chemistry and Biochemistry of the CAS) for help with the revision experiments. The work was supported by the European Regional Development Fund, OP RDE, Project: CARAT (No. CZ.02.1.01/0.0/0.0/16_026/0008382), Charles University project Progress Q43, and by the Czech Academy of Sciences – Strategy AV21 – Research program “VP29 Towards Precision Medicine and Gene Therapy.” Microscopy was performed in the Laboratory of Confocal and Fluorescence Microscopy co-financed by the European Regional Development Fund and the state budget of the Czech Republic, projects no. CZ.1.05/4.1.00/16.0347 and CZ.2.16/3.1.00/21515. The irradiation of HPHT nanodiamond by electrons was supported through Czech Academy of Sciences Project No. RVO61389005. We acknowledge the Electron Microscopy Core Facility, IMG ASCR, Prague, Czechia, supported by MEYS CR (LM2018129, LM2023050, CZ.02.1.01/0.0/0.0/18_046/0016045, CZ.02.1.01/0.0/0.0/16_013/0001775).

## Notes

We note that in some of our previous works on polymer coating of nanodiamonds,^12,39,76–82^ we used incorrect terminology. Although we obviously used the “grafting through” method in those studies, we referred to it as “grafting from.” In this article, the terminology is used correctly.

## Table of contents entry

Nanodiamonds coated with a random cationic copolymer consisting of (2-dimethylaminoethyl) methacrylate (DMAEMA) and N-(2-hydroxypropyl) methacrylamide (HPMA) monomers enable highly effective cellular delivery of siRNAs. Clathrin-mediated endocytosis as the predominant entry mechanism.

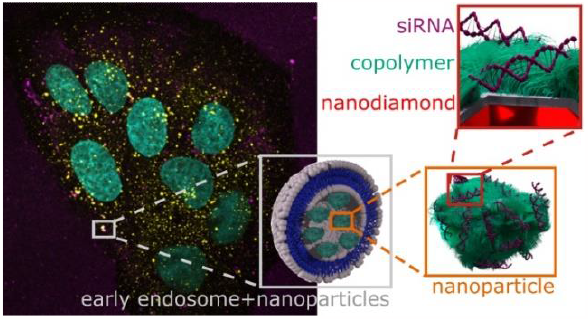

