## Supporting Information for "Cellular Uptake and Fate of Cationic Polymer-Coated Nanodiamonds Delivering siRNA: A Mechanistic Study"

Electronic Supporting Information

**Table S1.** Electrophoretic light scattering analysis of the $Cop^+$-FND and $Cop^+$-FND:siRNA complex in aqueous solution.

| | Apparent ζ-potential (mV) | Electrophoretic mobility (μm·cm/V·s) | Conductivity (μS/cm) | Applied voltage (V) |
|---|---|---|---|---|
| $Cop^+$-FND | 35.8 ± 3.9 | 2.81 ± 0.31 | 6.28 ± 0.15 | 9.97 |
| $Cop^+$-FND:siRNA* | 33.8 ± 0.5 | 2.65 ± 0.04 | 31.7 ± 0.5 | 4.96 |

All data represent the mean value ± standard deviation over four measurements.
*mass ratio 26:1

### Release of siRNA from $Cop^+$-FND:siRNA complex at different pH conditions

In comparison to the standard preparation procedure of $Cop^+$-FND:siRNA (see section **Complexation of $Cop^+$-FND:siRNA and Lipofect:siRNA**), the volume of water in the complexation reaction was reduced to obtain ~12× concentrated $Cop^+$-FND:siRNA sample without affecting its colloidal properties ($Cop^+$-FND:siRNA mass ratio was kept 26:1); 14.4 μL (26.8 mg/mL) $Cop^+$-FND + 10.6 μL (100 μM) siRNA. The final $Cop^+$-FND:siRNA complexes (15.4 mg $Cop^+$-FND/mL) were mixed with a buffer of defined pH value

[HEPES (10 mM), MES (10 mM), $NH_4OAc$ (10 mM), NaCl (130 mM)] resulting in 1:3 dilution of $Cop^+$-FND:siRNA.

Prior the mixing, the initial buffer solutions were adjusted at pH values: 4.0, 5.0, 6.0 and 7.0 (Metrohm pH microelectrode); the final volume of the samples for pH measurement was 100 μL. The $Cop^+$-FND:siRNA and control samples (see Table S2) were incubated for 30 min in the presence of buffer at RT, and pH was measured. To estimate the amount of free siRNA, the sample was quantitatively centrifuged for 20 min at 20,000 g and the supernatant was analyzed using Qubit™ miRNA assay kit. The Qubit assay allows the measurement of supernatant volume in the range of 1–20 μL, thereby increasing the quantification range (see the last column of Table S2). To get a high sensitivity of siRNA detection for nanoparticle-containing samples, 20 μL aliquots were analyzed. On the contrary, to reach an optimal quantification range for the positive control (free siRNA only), 1 μL was analyzed. To compare both types of measurements, Table S3 shows the recalculated siRNA concentration for positive control. Based on these calculations, the maximal siRNA concentration in each nanoparticle-containing sample (100% release of the siRNA considered) is ~15 times higher than the upper limit of quantification. Thus, the values <2.5 ng/mL in Table S2 indicate no significant release of siRNA from the surface of nanoparticles under the used conditions (less than 0.02 % of siRNA was released from the particle surface). These results also confirmed the effective complexation of the siRNA for the $Cop^+$-FND:siRNA mass ratio of 26:1 used throughout the study.

**Table S2.** Release of siRNA from $Cop^+$-FND:siRNA complex (mass ratio 26:1) at different pH conditions. For photographs of Cop+-FND:siRNA dispersions see **Figure S9**.

| | **Released amount of siRNA (ng/mL)*, ** /** pH | | | | Measured supernatant volume + Qubit assay volume (uL) |
|---|---|---|---|---|---|
| $Cop^+$-FND:siRNA | < **2.5** / 4.38 | < **2.5** / 5.31 | < **2.5** / 6.21 | < **2.5** / 7.05 | 20 + 180 |
| $Cop^+$-FND | < **2.5** / 4.65 | < **2.5** / 5.63 | < **2.5** / 6.47 | < **2.5** / 7.30 | 20 + 180 |
| siRNA (positive control)** | **542±1** / 3.76 | **549±1** / 4.66 | **583±1** / 5.65 | **580±1** / 6.68 | 1 + 199 |
| water (negative control) | < **2.5** / 3.96 | < **2.5** / 4.97 | < **2.5** / 5.99 | < **2.5** / 7.01 | 20 + 180 |

siRNA (MW = 13968, 100 μM)

*Assay tube concentration – siRNA concentration after dilution by Qubit assay; the effective quantification range for the assay tube concentration is 2.5–800 ng $mL^{-1}$

**All values were averaged over three measurements (technical replicates).

**Table S3.** Measured and theoretically calculated concentration of positive control from Table S2 for different volumes of analyzed supernatant.

| | | Released amount of siRNA (ng/mL)* | Measured supernatant volume + Qubit assay volume (uL) |
|---|---|---|---|
| siRNA (positive control) | measured | 580** | 1 + 199 |
| siRNA (positive control) | calculated*** | 740 | 1 + 199 |
| siRNA (positive control) | calculated*** | 14806 | 20 + 180 |
| **siRNA (positive control)** | **calculated****** | **11605** | **20 + 180** |

siRNA (MW = 13968, 100 μM)

*Assay tube concentration – siRNA concentration after dilution by Qubit assay

**Value from Table S2, pH 6.68

***Calculated siRNA concentration after dilution by Qubit assay if the true siRNA stock concentration is 100 μM

****Calculated and corrected concentration based on the known measured concentration from the first row.

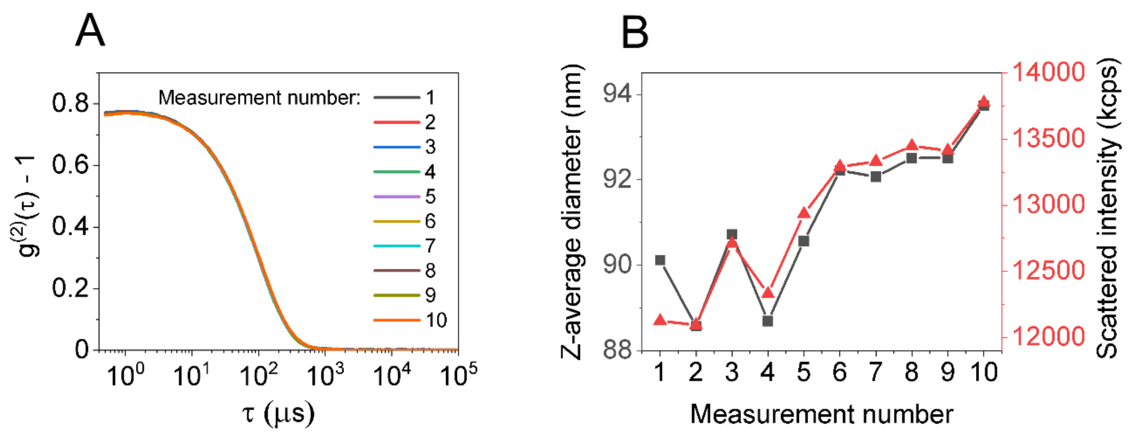


**Figure S1.** Stability testing of $Cop^{+}$-FND:siGAPDH complex (mass ratio 26:1) in full cell culture media at 37 °C. (A) DLS intensity autocorrelation functions. The measurement was repeated ten times (over approximately 30 min). (B) Scattered intensity (derived count rate) and Z-average size inferred from the $g^{(2)}(\tau)$–1 shown in panel A.

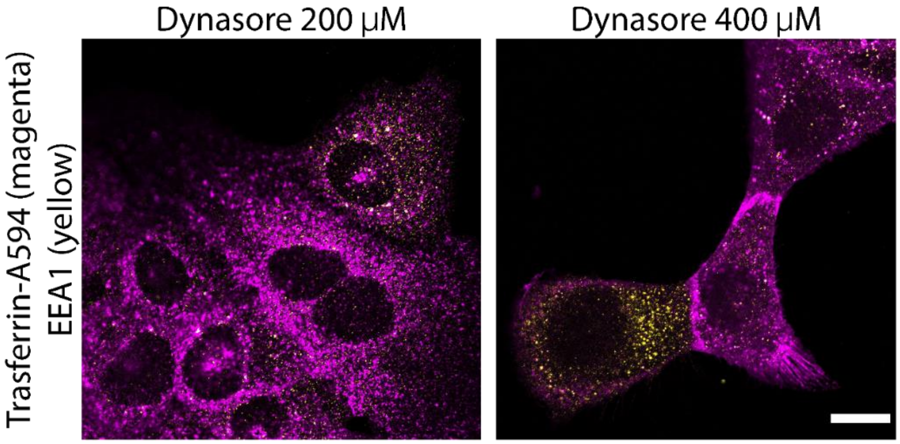


**Figure S2.** Determination of appropriate inhibition concentration for Dynasore. U-2 OS cells were pretreated in serum-free DMEM with Dynasore to final concentrations of 200 and 400 μM. After 20 min, cells were chilled (3 min, 7 °C). Transferrin-A594 (25 μg/mL) was pipetted to sample wells. Cells were washed with PBS and fixed with paraformaldehyde after 55 min. 400 μM inhibitor concentration resulted in decreased amount of punctate transferrin signal in cytoplasm. Scale bar = 20 μm.

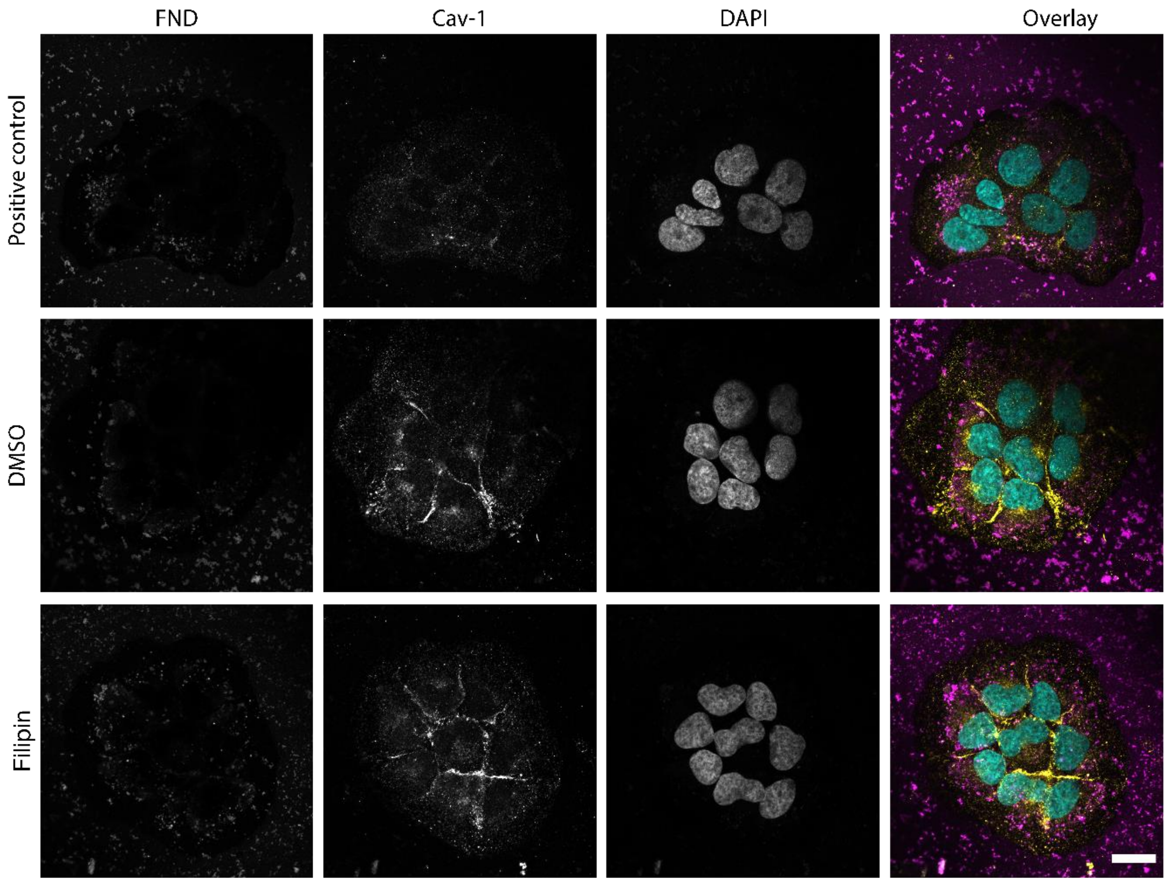


**Figure S3.** Effect of filipin on the uptake of $Cop^+$-FND:siGAPDH complex. U-2 OS cells were pretreated in serum-free DMEM (positive control), with filipin (5 µg/mL in DMSO) or DMSO. After 20 min, cells were chilled (3 min, 7 °C). $Cop^+$-FND:siGAPDH complex was pipetted to sample wells. Cells were washed with PBS and fixed with paraformaldehyde after 55 min. Amount of FND signal within the cytoplasm suggested negligible effect of filipin on the uptake of FNDs. Scale bar = 20 µm.

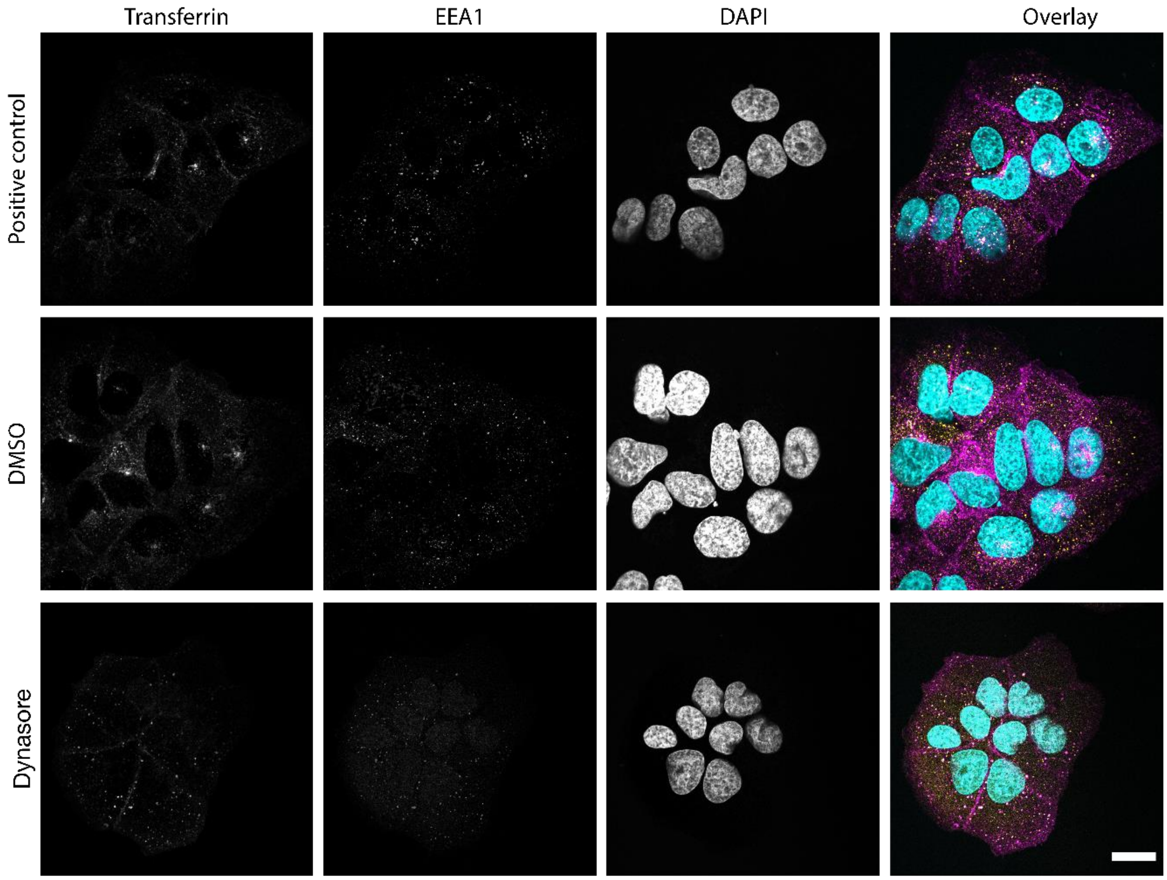


**Figure S4.** Validation of appropriate inhibition concentration for Dynasore. U-2 OS cells were pretreated in serum-free DMEM, with Dynasore (400 μM) or DMSO. After 20 min, cells were chilled (3 min, 7 °C). Transferrin-A594 (25 μg/mL) was pipetted to sample wells. Cells were washed with PBS and fixed with paraformaldehyde after 55 min. When compared with the positive control and DMSO-treated cells, the cytoplasmic Transferrin-A594 signal in Dynasore-treated cells was significantly reduced. Scale bar = 20 μm.

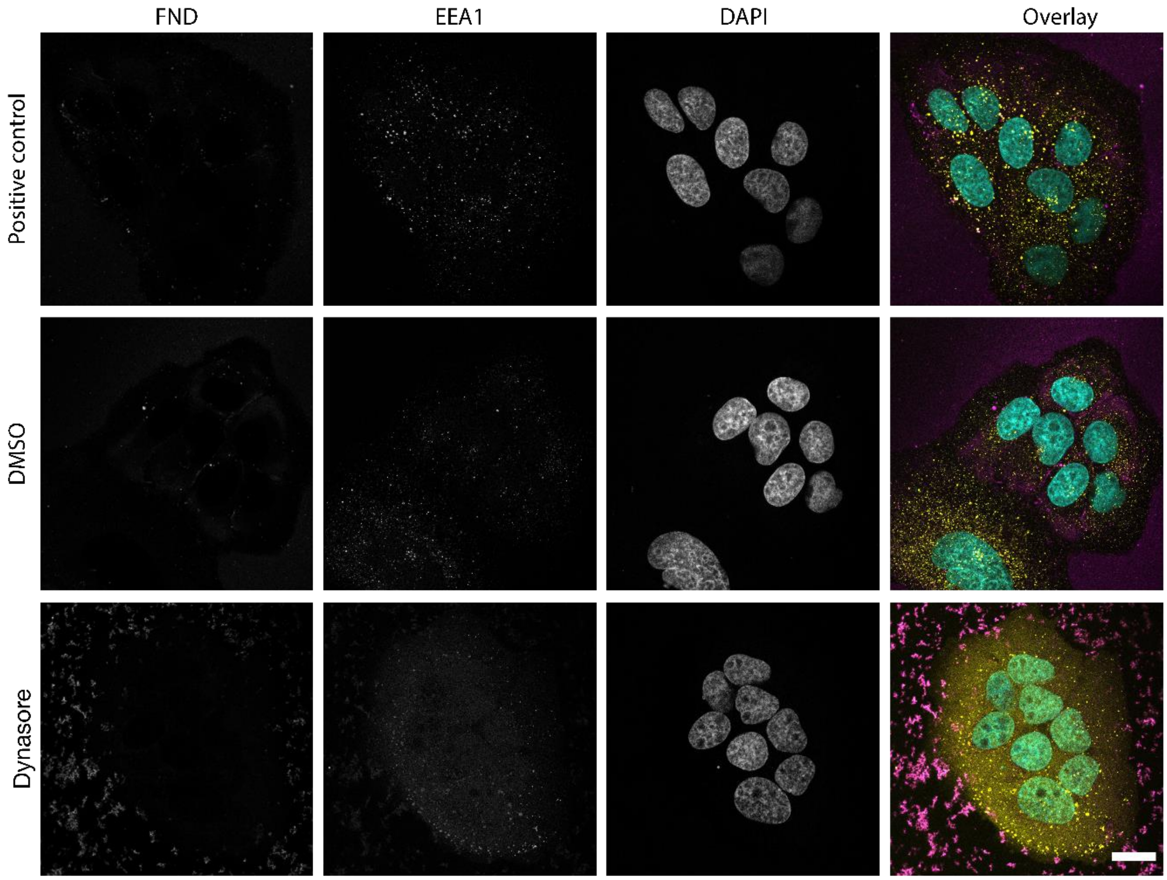


**Figure S5.** Effect of Dynasore on the uptake of Cop$^+$-FND:siGAPDH complex. U-2 OS cells were pretreated in serum-free DMEM, Dynasore (400 μM), and DMSO. After 20 min, cells were chilled (3 min, 7 °C). Cop$^+$-FND:siGAPDH complex was pipetted to sample wells. Cells were washed with PBS and fixed with paraformaldehyde after 55 min. When compared with the positive control and DMSO-treated cells, the cytoplasmic FND signal in Dynasore-treated cells was absent. Scale bar = 20 μm.

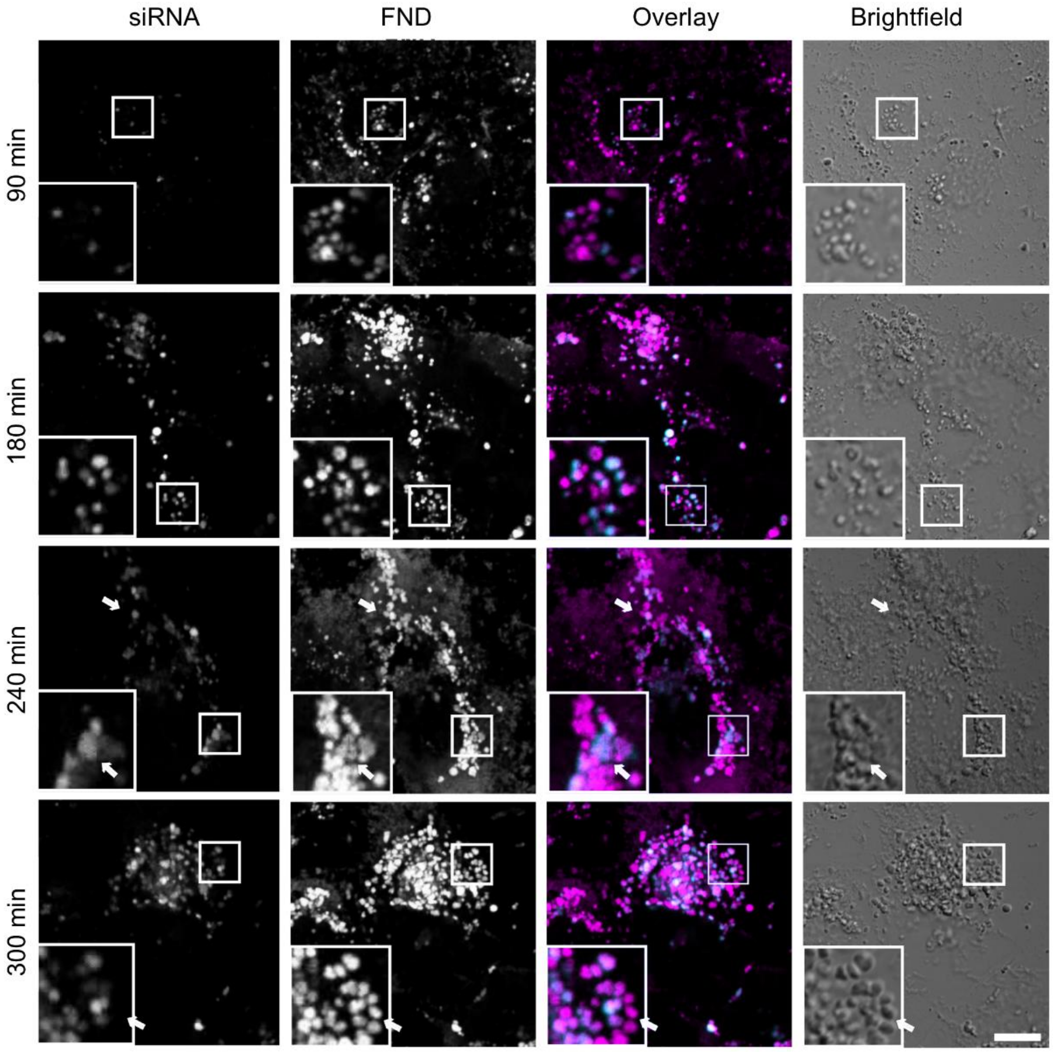


**Figure S6.** Temporal visualization of accumulation and spatial distribution of Cop+-FND:siGAPDH complexes (FND magenta, siGAPDH cyan) within fixed U-2 OS cells using fluorescence microscopy. The cells were incubated with the FND-siGAPDH complex (140 nM) until fixed with paraformaldehyde at timepoints up to 6 hours. Arrows indicate heterogeneities in spatial overlap of the FND and siGAPDH signal. The scale bar represents 20 μm.

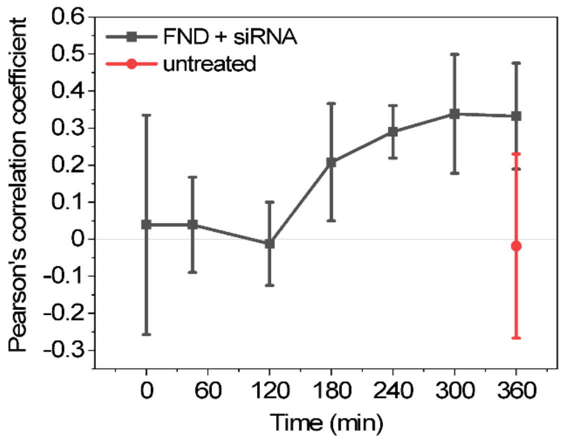


**Figure S7.** Colocalization analysis of FND and siRNA (GAPDH) expressed by Pearson's correlation coefficient ($n$ = 3 experiments).

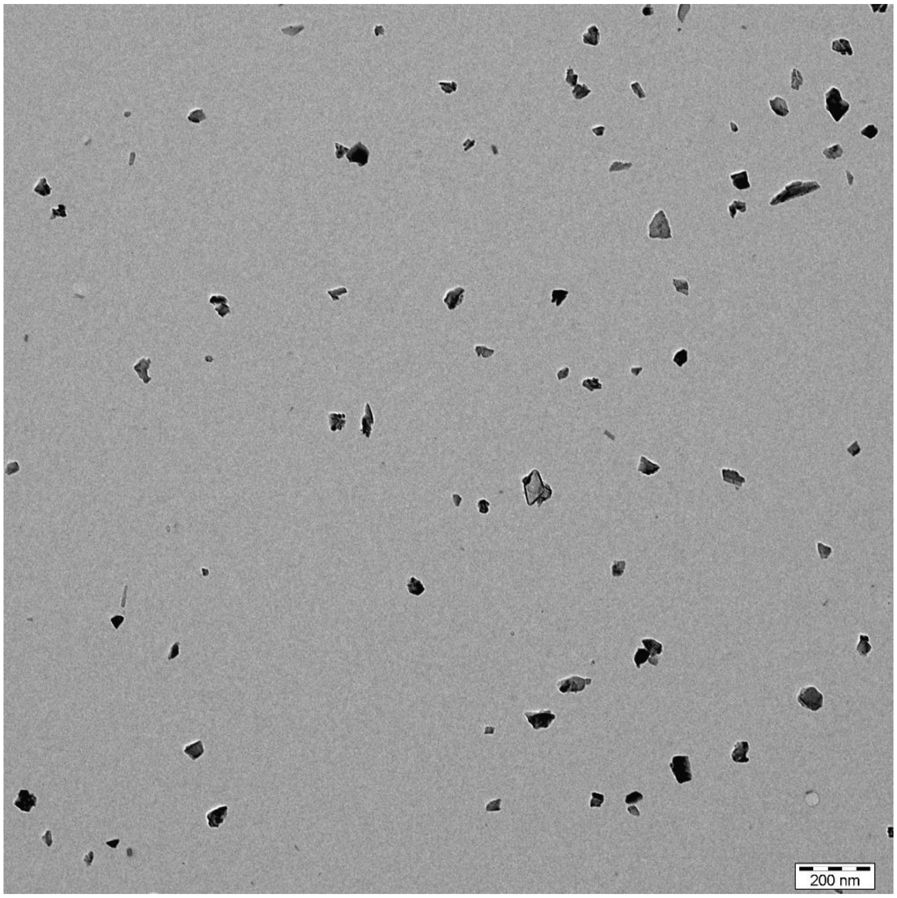


**Figure S8.** TEM image of $Cop^+$-FND sample. Scale bar = 200 nm.

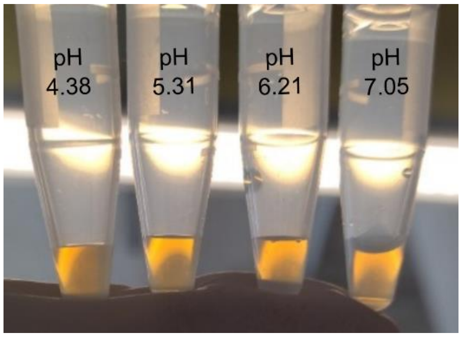


**Figure S9.** Photograph of $Cop^+$-FND:siRNA dispersions (3.9 mg/mL) at various pH conditions after 30 min at RT.

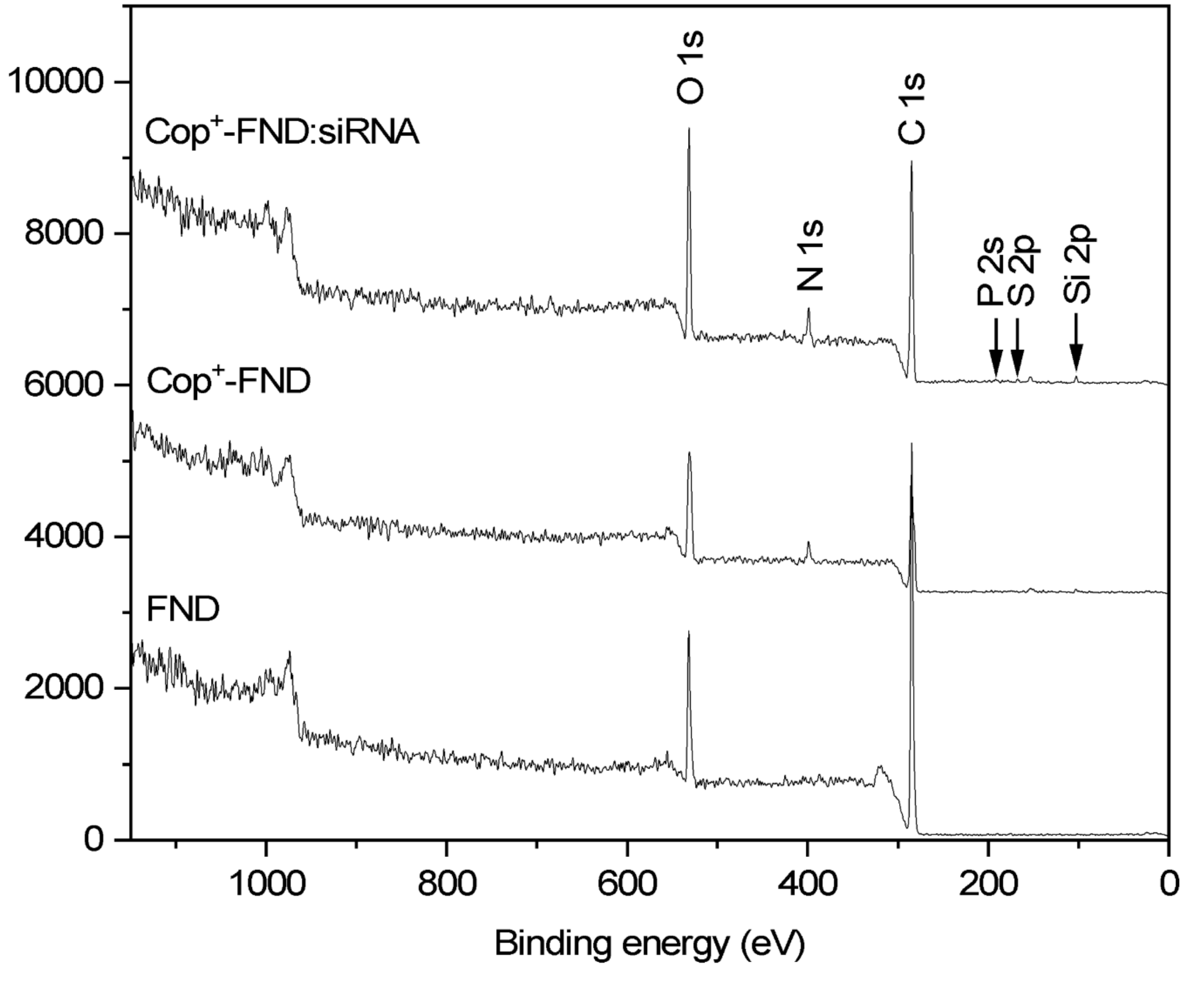


**Figure S10.** XPS spectra of the starting material (FND), $Cop^+$-FND and its complex with siRNA ($Cop^+$-FND:siRNA). The peaks of interest are individually marked. Curves are offset for clarity, but the intensity scale is identical for all spectra.

**Table S4**. Elemental composition (atomic %) obtained from XPS analysis of FND, $Cop^{+}$-FND and $Cop^{+}$-FND:siRNA.

| | C | O | N | Si | P | S |
|---|---|---|---|---|---|---|
| FND | 89.5 | 10.3 | 0.0 | 0.22 | — | — |
| $Cop^{+}$-FND | 75.7 | 18.4 | 3.5 | 2.41 | — | — |
| $Cop^{+}$-FND:siRNA | 73.7 | 18.4 | 5.6 | 1.92 | 0.21 | 0.17 |
